# Metaproteome analysis of short-term thermal stress in three sympatric coral species reveals divergent host responses

**DOI:** 10.1101/2025.03.25.645042

**Authors:** Shrinivas Nandi, Timothy G. Stephens, Erin E. Chille, Samantha Goyen, Line K. Bay, Debashish Bhattacharya

## Abstract

The accelerating loss of coral reefs worldwide due to anthropogenic climate change has led to a myriad of studies aimed at understanding the basis of coral resilience to support reef conservation. Here, we integrate physiological measurements with proteomic and metabolomic profiles to examine species-specific responses to increased temperature in three sympatric reef-building corals from the Great Barrier Reef: *Acropora hyacinthus*, *Porites lobata*, and *Stylophora pistillata*. We find species-specific stress response strategies with *A. hyacinthus*, a thermally sensitive species, exhibiting rapid decline in endosymbiont physiology, coupled with a one-third reduction in protein abundance. In contrast, *P. lobata* displayed a delayed physiological response to stress and a muted proteome response, suggesting greater resilience. *S. pistillata* initially showed minor shifts in the proteome followed by colony “bail-out”. Overall, we observed markedly different responses in most biochemical pathways in the three coral species. Nonetheless, some known biomarkers of stress, including heat-shock proteins, showed conserved responses to thermal stress with differences in temporal abundance reflecting bleaching resistance. Our results underscore the species-specific nature of coral responses to thermal stress and highlight proteomic signatures associated with symbiosis breakdown, offering mechanistic insights into coral bleaching susceptibility and resilience.

## Introduction

Stony corals (Scleractinia) thrive in oligotrophic waters due to their ability to form a symbiotic association with dinoflagellate algae in the family Symbiodiniaceae (Gordon and Leggat, 2010). These photosynthetic endosymbionts sustain the host coral through the transfer of photosynthates (Muscatine and Porter, 1977). In exchange, corals provide Symbiodiniaceae with a stable, sheltered environment, inorganic carbon, and other essential nutrients (Frankowiak et al., 2016). Algal symbionts are contained within host gastrodermal cells in symbiosome compartments which provide an intracellular microenvironment optimized for photosynthesis (Barott et al., 2022, 2015; Bertucci et al., 2013; Thies et al., 2022). Under stressful conditions (e.g., high nutrient enrichment, cold- and heat-stress), coral hosts may lose their endosymbiotic algae, which is known as coral bleaching. During bleaching, some corals can sustain themselves for extended periods of time through heterotrophic feeding (Grottoli et al., 2006). However, bleached corals are more susceptible to disease and may succumb to starvation if the symbiosis is not reestablished (Bourne et al., 2016; Muller et al., 2008). The increase in mean global oceanic temperature, a result of anthropogenic climate change, has led to increased bleaching episodes and a significant decline in global coral populations (Hughes et al., 2017).

Although coral bleaching is a conserved stress response across Scleractinia, bleaching risk and the degree of bleaching for a colony depends on several factors. Coral bleaching responses are shaped by a complex interplay of ecological, physiological, and evolutionary factors, leading to species-specific differences in susceptibility (Loya et al., 2001; Woesik et al., 2011). Bleaching outcomes under different levels of thermal stress have been extensively studied in the Great Barrier Reef, Australia (Burn et al., 2023; Hughes et al., 2019, 2018, 2017; Pratchett et al., 2020). For instance, massive and slow-growing corals such as *Porites* often exhibit greater thermal tolerance when compared to branching, fast-growing species such as *Acropora*, which tend to bleach more readily (Burn et al., 2023; Fitt et al., 2009; Marshall and Baird, 2000; Pratchett et al., 2020), likely reflecting different strategies for energy allocation (i.e., resilience vs. rapid growth) of these long-lived sessile organisms under dyanamic environmental conditions. Variations in skeletal morphology (Swain et al., 2018, 2016), tissue thickness (Barkley et al., 2018; Qin et al., 2020), and metabolic strategies, such as heterotrophic plasticity (Grottoli et al., 2006; Martinez et al., 2024; Sangmanee et al., 2020), can contribute to differential tolerance to thermal and other environmental stressors.

Variations in algal symbiont composition have been extensively studied, with some algal lineages conferring higher thermal tolerance to the coral colony (Bhattacharya et al., 2024; Buerger et al., 2020; Putnam et al., 2017; Rowan, 2004; Wang et al., 2023; Ziegler et al., 2018). The prokaryotic microbiome may also play a major role in coral holobiont resilience (Peixoto et al., 2017; Reshef et al., 2006; Rosado et al., 2019; Ziegler et al., 2017). In addition, local environmental conditions including water flow, light intensity, nutrient levels, and previous exposure to stress modulate bleaching susceptibility. Corals in high-flow or turbid environments may experience stress buffering, whereas those inhabiting sheltered, low-flow conditions are more likely to experience thermal extremes (Morgan et al., 2017; Risk and Edinger, 2011). Prior exposure to sublethal stress, or high frequency thermal variability can induce acclimatization or select for stress-tolerant genotypes, resulting in site- and species-specific patterns in bleaching severity (Barshis et al., 2013; Kenkel et al., 2013; Oliver and Palumbi, 2011; Safaie et al., 2018).

Developing reliable molecular biomarkers to detect early stress responses in corals is a major goal in reef monitoring and conservation. These tools will enable rapid, pre-bleaching health assessment which can guide targeted interventions (Chille et al., 2025; Downs et al., 2005). Transcriptomic datasets have been extensively used to identify putative biomarkers of thermal stress (Chille et al., 2025; Cowen and Putnam, 2022; Cziesielski et al., 2019).

However, a central challenge is that stress produces high variation in gene expression levels in different species (Da-Anoy et al., 2024; Ip et al., 2022; Rosic et al., 2020), and even among genotypes within a species (Chille et al., 2024; Kenkel and Matz, 2016). Nonetheless, transcriptomic studies have identified some marker genes of thermal stress, including those encoding heat-shock proteins (e.g., HSP70, HSP90) (Nakamura et al., 2011; Rosic et al., 2011; Seveso et al., 2016), ubiquitin-related proteins (DeSalvo et al., 2008; Rodriguez-Lanetty et al., 2010, 2009; Seneca and Palumbi, 2015), and antioxidant enzymes (Downs et al., 2002; Krueger et al., 2015) among others. A recent review of 307 gene and protein expression papers studying stress response in cnidarians found that most genes have inconsistent responses across different experiments, with only 14 having shared responses across at least two species tested and NF-κB having shared responses across four species, thereby suggesting a high degree of species specificity vis-à-vis the coral stress response (Molinari et al., 2025).

The use of proteomics in coral research is relatively under-developed and may enhance biomarker identification by providing a more direct measure of the functional molecules that drive cellular processes, capturing post-transcriptional regulation and protein modification that transcriptomics alone cannot address (Greenbaum et al., 2003). In addition, protein biomarkers offer an advantage to mRNA because they are more readily integrated into existing point-of care diagnostic tools (Chille et al., 2025). Research across diverse eukaryotic systems, including the coral *Montipora capitata*, demonstrates that transcriptomic data often poorly represent the proteome, with weak correlations observed between changes in transcript abundance and accumulation of the corresponding proteins (Williams et al., 2023). Thus, proteomic data can provide a rich resource not only for biomarker development, but also for further elucidation of coral thermal stress responses (Chille et al., 2025).

Metabolomics provides complementary insights to proteomic data by capturing the end products of cellular activity, i.e., metabolites that directly reflect the real-time physiological state of the organism (Fiehn, 2002; Roessner and Bowne, 2009). Previous metabolomic studies in corals have reported changes in antioxidant-related metabolites during thermal stress (Williams et al., 2021b) and shifts in amino acid pools (Chiles et al., 2022; Martinez et al., 2020, 2022a). Notably, recent findings have demonstrated the accumulation of specific dipeptides as reliable indicators of pre-bleaching thermal stress. These include lysine-glutamine (KQ), arginine-glutamine (RQ), arginine-alanine (RA), and arginine-valine (RV) (Huffmyer et al., 2024; Williams et al., 2021a).

In this study, we used proteomic data to investigate the coral holobiont stress response in three sympatric species from the Great Barrier Reef (GBR), Australia: *Acropora hyacinthus*, *Porites lobata*, and *Stylophora pistillata* [**Figure 1**]. These species were selected due to their known differences in thermal tolerance: i.e., sensitive, resilient, and low-to-moderate sensitivity, respectively (Burn et al., 2023; Fitt et al., 2009; Manalili et al., 2025; Marshall and Baird, 2000; Pratchett et al., 2020). We examined key metabolic pathways in each species to better understand their responses to thermal stress and searched for conserved proteins across species that can potentially be used as biomarkers. Lastly, we investigated the metabolomic data to validate previously reported shifts in metabolite abundance, including dipeptides, amino acids, and antioxidants that play key roles in the coral thermal stress response.

**Figure 1.**
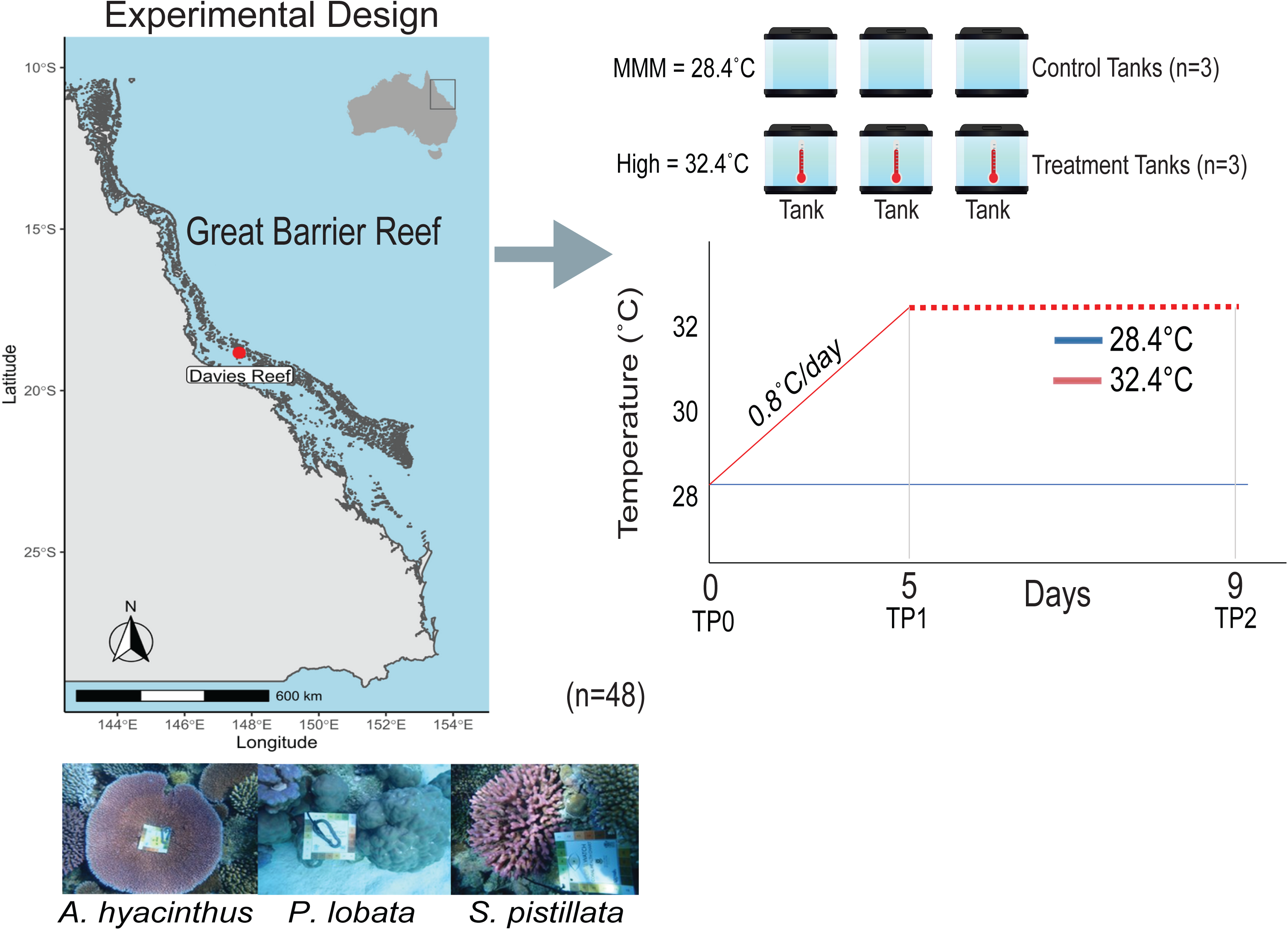
Study sites and experimental designs. Location of samples (*n* = 48) collected and experimental design (right panel) are shown. The coral images were generated by Samantha Goyen, map created in R using the Environmental Systems Research Institute (ESRI) shapefiles and Mapdata package, experimental setup image made in Canva. TP refers to the timepoint at which corals were sampled. Tank set up is displayed on the left with the mean temperatures of each tank shown. Graph showing the ramp up protocol used for the tanks, red, thermal stress tanks and blue, control tanks.

## Results

### Host bailout and mortality

No mortality was observed in *A. hyacinthus* and *P. lobata*. In contrast, *S. pistillata* exhibited significant tissue loss (i.e. “bail-out”) and loss of color (becoming paler) based on visual assessment at TP1 [**Supplemental Figure 1**] and complete tissue loss was observed by TP2, in most fragments.

### Endosymbiont community composition and physiology

Because coral bleaching is associated with reduced algal symbiont abundance in host tissues and reduced photosynthetic efficiency, cell counts [**Figure 2A**] and Pulse Amplitude Modulation (PAM) fluorometer measurements [**Figure 2B**] were taken for each of the samples collected at each time point to characterize the effect of thermal stress on endosymbiont physiology in the three coral species [**Supplemental Table S1a**]. Symbiont cell densities, calculated as a factor of coral tissue area; hereinafter, endosymbiont abundance, were used to assess bleaching severity and to normalize the proteomic data generated from the symbionts in each of the sampled coral fragments. The latter was done to account for reduced symbiont protein abundance values because of symbiont cell loss and not reduced accumulation of the protein within the symbiont cells. Under ambient conditions, endosymbiont abundances remained stable across the experiment [**Figure 2A**]. Using ANOVA, we observed significant shifts in endosymbiont abundance upon exposure to thermal stress in *A. hyacinthus* (df = 5, F = 6.032, *p-*value = 0.0051), but not in *P. lobata* (df = 5, F = 1.015, *p-*value = 0.450) or *S. pistillata* (df = 3, F = 3.285, *p-*value = 0.079). In *A. hyacinthus*, when comparing endosymbiont abundance between ambient conditions (TP1: 20.50 ± 0.32, TP2: 20.24 ± 0.52) and thermal stress samples (TP1: 19.23 ± 0.70, TP2: 18.36 ± 0.31), Tukey-HSD post hoc analysis indicated a significant difference only at TP2 (adj-*p*- value = 0.014).

**Figure 2.**
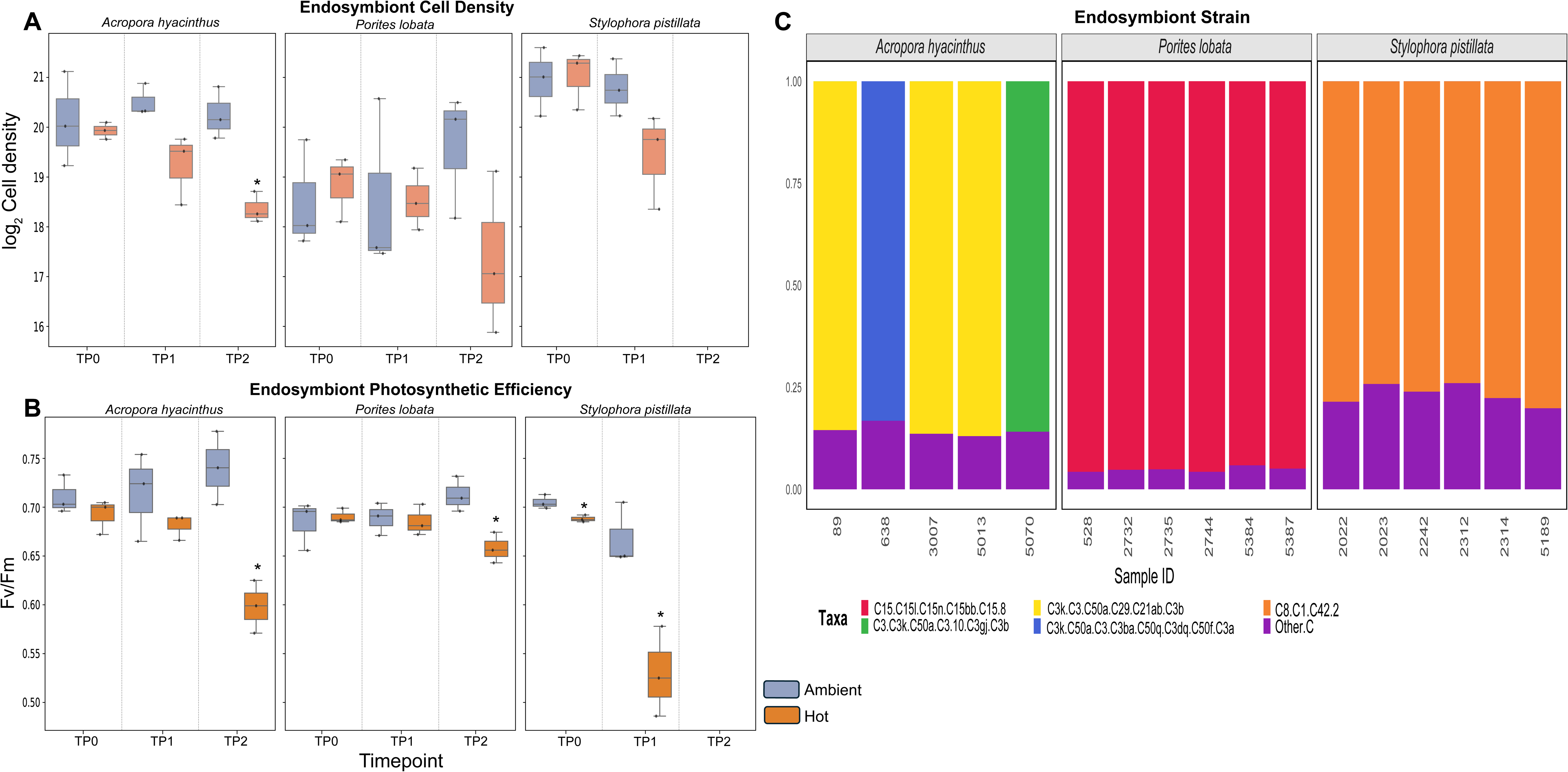
Physiological measurements of GBR corals under ambient and heat stress treatments. (**A**) Endosymbiont cell abundance (log_2_ transformed) over the duration of the experiment. Blue boxes represent ambient cell density at each timepoint (x-axis) and the orange boxes represent cell density upon exposure to thermal stress. The three graphs are the different species used in this study. The y-axis shows the log_2_ normalized cell density (cells/ml/cm^2^). **(B)** The y-axis shows the *F_v_/F_m_*ratio, which is a proxy for photosynthetic efficiency, and the x-axis represents the experimental time points. Blue boxes are the values for the ambient samples and the orange for the stress samples. * Represents statistical significance (*p*-value < 0.05) between the ambient and heat stress treatments at that timepoint. **(C)** Variation in major SymPortal ITS2 Symbiodiniaceae communities within *Acropora hyacinthus*, *Porites lobata*, and *Stylophora pistillata* colonies from the Great Barrier Reef. The y-axis is the relative abundance. One replicate of *P. lobata* consisted of <1% relative abundance of *Gerakladium* and has not been included in the image.

Same as endosymbiont abundance, photosynthetic efficiency (*F_v_/F_m_*) values remained stable under ambient conditions in all three species. However, under thermal stress, we observed significant declines in endosymbiont photosynthetic efficiency in all three coral species [**Figure 2B**]. ANOVA analysis showed significant photosynthetic efficiency shifts in *A. hyacinthus* (df = 5, F = 8.552, *p*-value = 0.0011). Tukey-HSD post hoc analyses revealed a significant decrease in thermal stress samples at TP2 (adj-*p*-value = 0.0007, Ambient = 0.740 ± 0.037, Hot = 0.59 ± 0.027). Similarly, in *P. lobata* (ANOVA: df = 5.0, F = 3.067, *p*-value = 0.051), a significant decline was observed only at TP2 (adj-*p*-value = 0.020, Ambient = 0.712 ± 0.018, Hot = 0.657 ± 0.015). Lastly, in *S. pistillata* (df = 5, F = 23.89, *p*-value = 0.00024), we observed a significant decline at TP1 (adj-*p*-value = 0.0015, Ambient = 0.668 ± 0.032, Hot = 0.529 ± 0.046).

We also characterized the endosymbiont community profiles using ITS2 sequencing [**Figure 2C**] to determine whether differences in host performance could be attributed to differences in resident endosymbiont composition. ITS2-based *Symbiodiniaceae* profiling of the three coral species using SymPortal showed that all samples associated exclusively with the genus *Cladocopium* (except for one *P. lobata* replicate, which had <1% relative abundance of *Gerakladium*) [**Figure 2C**]. The three coral species exhibited distinct major ITS2 type profiles, which were C15-C15l-C15n-C15bb-C15.8 for all samples of *P. lobata*, C8/C1- C42.2 for all samples of *S. pistillata*, and C3k/C3-C50a-C29-C21ab-C3b, C3/C3k-C50a-C3.10-C3gj-C3b, and C3k-C50a-C3-C3ba-C50q-C3dq-C50f-C3a for all samples of *A. hyacinthus* [**Supplemental Table 1b**].

### Overall host proteome analysis

Prior to filtering, we obtained 3,793 protein groups (hereinafter proteins) from *A. hyacinthus* **[Supplemental Table 2]**, 4,098 proteins from *P. lobata* **[Supplemental Table 3]**, and 2,479 proteins from *S. pistillata* **[Supplemental Table 4]**. After filtering for unique and razor peptides and removing proteins with low counts, 2,255 (*A. hyacinthus*), 2,351 (*P. lobata*), and 1,335 (*S. pistillata*) proteins remained for downstream analysis. We assessed differential abundance for all proteins using pairwise comparisons. A protein was considered to have increased abundance if it had a log_2_ fold-change (FC) > 0.5 and *p*-value < 0.05, and decreased abundance if it had a FC < –0.5 with *p*-value < 0.05.

Differentially abundant proteins were detected at TP0 in all three species, suggesting the presence of tank effects. In *A. hyacinthus*, 72 proteins showed increased abundance and 5 showed decreased abundance **[Supplemental Figure 2]**. In *P. lobata*, 46 proteins had increased abundance and 82 had decreased abundance. In *S. pistillata*, 5 proteins exhibited increased abundance and 148 exhibited decreased abundance. Although these results indicated tank effects at TP0, we observed a markedly higher number of differentially abundant proteins at later timepoints under thermal stress. In *A. hyacinthus* at TP1, 23 proteins showed increased abundance and 161 showed decreased abundance; at TP2, 75 proteins showed increased abundance and 101 showed decreased abundance. In *P. lobata* at TP1, 23 proteins showed increased abundance and 96 showed decreased abundance; at TP2, 86 proteins showed increased abundance and 235 showed decreased abundance **[Supplemental Figure 2]**. Lastly, in *S. pistillata*, 87 proteins showed increased abundance and 57 showed decreased abundance at TP1. Samples were collected only for two timepoints, due to complete bailout by TP2 (**see Discussion**).

### Pathway analysis of coral hosts

Enrichment analysis at the “B-level” (KEGG Orthology) was done to compare pathway-level differences in coral stress responses. Pathways were considered upregulated if ≥5% of the total proteins in a pathway showed increased abundance under stress, and downregulated if ≥5% showed a decrease. Mixed responses were assigned when both patterns were observed. After filtering for pathways with more 10 proteins in each species, 25 pathways were retained for analysis [**Figure 3A**]. At TP1, *A. hyacinthus* showed downregulation in 16 pathways; *P. lobata* showed downregulation in 10 pathways and a mixed response in one; *S. pistillata* showed upregulation in seven pathways, downregulation in three, and a mixed response in three. By TP2, *A. hyacinthus* displayed a shift, with 5 pathways downregulated, six upregulated, and one mixed. In contrast, *P. lobata* exhibited stronger downregulation at TP2, with 16 pathways downregulated, five mixed, and one upregulated. Several pathways downregulated in *P. lobata* at TP2 mirrored those observed in *A. hyacinthus* at TP1 (e.g., energy metabolism).

**Figure 3.**
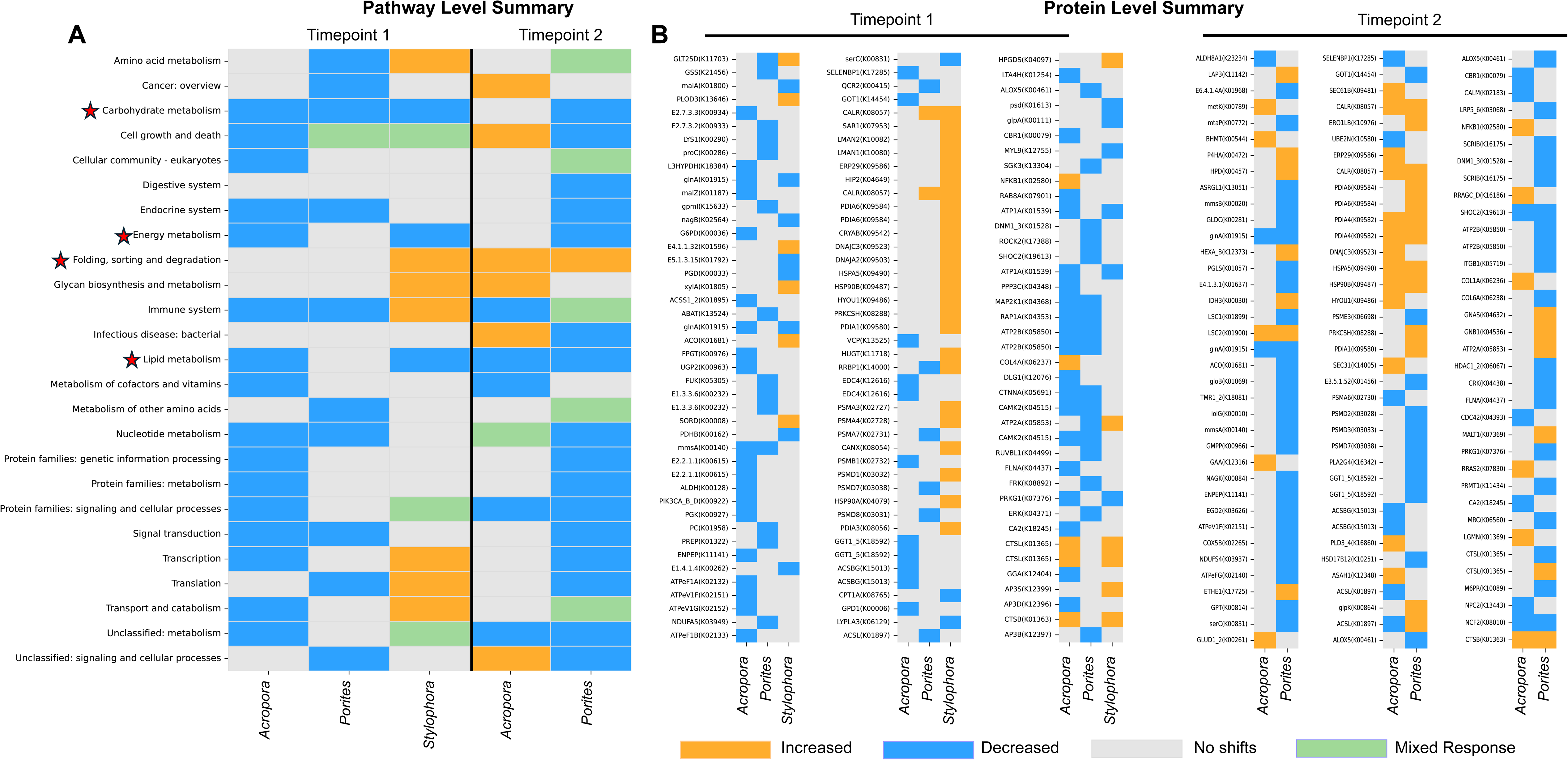
Proteomic responses at the pathway level and individual protein abundances in conserved pathways. **(A)** Heatmap showing the pathway level response (y-axis) for each coral species (x-axis) at each timepoint. The blue boxes are downregulated pathways in a species at that timepoint, the orange boxes show upregulated pathways, green boxes show a mixed response, and grey boxes indicate no change (see figure legend). The red stars indicate conserved pathways that are extensively discussed in the main text. **(B)** Heatmap showing significant protein responses (|FC| > 0.5 and *p*-value < 0.05), with proteins on the y-axis (Name and KEGG ID) and the species on the x-axis. The graph is divided in two timepoints, TP1 left) and TP2 (right). Some proteins may be listed twice; this is due to the presence of multiple copies in a species that are significant.

To identify shared vs. species-specific responses, we examined conserved patterns, defined as a consistent direction of regulation across all species, and divergent responses (those not conserved). Pathways were considered conserved even if their trends emerged at different timepoints. For example, energy metabolism was downregulated at TP1 in both *S. pistillata* and *A. hyacinthus*, but in *P. lobata*, this downregulation occurred only at TP2. This approach accounts for differences in bleaching rates among the species, as evident from physiological measurements. Only four pathways showed conserved response wherein carbohydrate metabolism, energy metabolism, lipid metabolism were downregulated, and folding/sorting/degradation was upregulated [**Figure 3A, red stars].** In contrast, 21 pathways displayed divergent responses (for example: cell growth and death) or species-specific responses (for example: downregulation of “digestive system” in *P. lobata* only).

We prioritized conserved pathways for further functional analysis. Although “amino acid metabolism” showed divergent trends, it was included due to its known role in thermal stress in corals. Furthermore, two custom sets of proteins, symbiosome maintenance (lysosome, phagosome proteins, carbonic anhydrases, and Vtype ATPases) and reactive oxygen species (ROS) detoxification enzymes were assessed due to their known role in coral bleaching (**Supplemental Table 5, See Supplemental Text**). Some proteins may be part of multiple pathways but are presented only once in the results. **Figure 3B** summarizes the results presented below.

### Shared proteome responses in coral hosts

#### Carbohydrate metabolism

All three coral species exhibited downregulation of carbohydrate metabolism under thermal stress, with 39 proteins differentially abundant. However, only two proteins were shared across species, although they were differentially abundant at different timepoints. Malonate-semialdehyde dehydrogenase (ALDH6A1, K00140), involved in propionate metabolism, showed decreased abundance at TP1 in both *A. hyacinthus* (FC = –0.52, *p*-value = 0.0086) and *P. lobata* (FC = –0.72, *p*-value = 0.011), and at TP2 in *P. lobata* (FC = –0.77, *p*-value = 0.025). Aconitate hydratase (ACO, K01681) exhibited a divergent pattern, with increased abundance in *S. pistillata* at TP1 (FC = 1.09, *p*-value = 0.032) and decreased abundance in *P. lobata* at TP2 (FC = –0.58, *p*-value = 0.044). The remaining 37 proteins were species-specific **[Figure 3B]**.

#### Energy metabolism

Downregulation of energy metabolism was observed in all three species. Key proteins in oxidative phosphorylation pathways, including ATPases, NADHs, and cytochrome proteins, were differentially abundant. In *A. hyacinthus* at TP1, ATPeF1A (K02132, FC = –0.50, *p*- value = 0.037) and ATPeF1B (K02133, FC = –0.53, *p*-value = 0.035) both showed decreased abundance. Additionally, the V-type ATPase (VHA) genes ATPeV1G (K02152, FC = –1.57, *p*-value = 0.027) and ATPeV1F (K02151, FC = –1.44, *p*-value = 0.031) also showed decreased abundance at TP1. None of these proteins were statistically significant at TP2.

Different proteins related to energy metabolism were differentially abundant in *P. lobata*. At TP1, NADH dehydrogenase subunit NDUFA5 (K03949, FC = –0.60, *p*-value = 0.035) showed decreased abundance. A stronger trend was observed in the same pathway at TP2, wherein ATPeFG (K02140, FC = –0.79, *p*-value = 0.042), NDUFS4 (K03937, FC = –0.67, *p*-value = 0.010), and ATPeV1F (FC = –1.49, *p*-value = 0.041) all showed decreased abundance in *P. lobata*. No significant shifts in oxidative phosphorylation proteins were observed in *S. pistillata*.

#### Amino acid metabolism

Amino acids play an essential role in the coral-algal symbiosis and are catabolized by the host under thermal stress. Glutamine synthetase (GLNA; K01915) showed decreased abundance in *A. hyacinthus* at both TP1 (FC = –0.98, *p*-value = 0.028) and TP2 (FC = –1.09, *p*-value = 0.020). In *S. pistillata*, two copies of GLNA were detected, both of which showed decreased abundance at TP1 (FC = –0.64, *p*-value = 0.0043 and FC = –0.74, *p*-value = 0.00032). No shifts in GLNA abundance were observed in *P. lobata* at TP1, however, a significant decrease was observed at TP2 (FC = –2.35, *p*-value = 0.0051). We also assessed the abundance of glutamate dehydrogenase (GDH). One GDH protein (K00262) was detected in *S. pistillata*, which showed a significant decrease at TP1 (FC = –0.50, *p*-value = 0.0029).

In *A. hyacinthus*, a different GDH protein (K00261) was identified that showed increased abundance at TP2 (FC = 0.80, *p*-value = 0.016). In *P. lobata*, no significant shifts in GDH abundance were detected. Lastly, we assessed 4-hydroxyphenylpyruvate dioxygenase (HPPD), which showed a significant increase only in *P. lobata* at TP2 (K00457, FC = 0.68, *p*-value = 0.028).

#### Protein folding, sorting, and degradation

Elevation of heat shock protein (HSP) abundance is one of most conserved responses in corals under thermal stress (Molinari et al., 2025) and showed significant responses in all three species. At TP1, three HSPs showed a significant increase in *S. pistillata* only: HSPA5 (K09490, FC = 1.16, *p*-value = 0.0013), HSP90B (K09487, FC = 0.96, *p*-value = 0.00057),

and HSP90A (K04079, FC = 0.78, *p*-value = 0.007). At TP2, significant increases were observed in both *A. hyacinthus* and *P. lobata* for HSPA5 (FC = 0.60, *p*-value = 0.016; FC = 0.57, *p*-value = 0.028, respectively) and HSP90B (FC = 1.01, *p*-value = 0.00086; FC = 1.26, *p*-value = 0.0022). Additionally, DNAJ proteins (HSP40 family) showed similar patterns: at TP1, two proteins increased in *S. pistillata*, DNAJC3 (K09523, FC = 1.22, *p*-value = 0.00083) and DNAJA2 (K09503, FC = 2.05, *p*-value = 0.010). At TP2, *A. hyacinthus* also showed increased abundance of DNAJC3 (FC = 1.07, *p*-value = 0.016). Another chaperone protein, Calreticulin (CALR, K08057), showed a staggered yet consistent response. At TP1, it showed increased abundance in *P. lobata* (FC = 0.71, *p*-value = 0.026) and *S. pistillata* (FC = 0.89, *p*-value = 0.0065). At TP2, we observed a significant increase in both *A. hyacinthus* (FC = 0.70, *p*-value = 0.0058) and *P. lobata* (FC = 1.02, *p*-value = 0.0081). We also observed significant shifts in proteasome subunit proteins involved in degradation **[See Supplemental Text]**.

#### Lipid metabolism

Shifts in lipid content are known markers of coral thermal stress. We observed two key fatty acid biosynthesis genes that showed conserved and significant responses to thermal stress: long-chain-fatty-acid--CoA ligase (ACSBG, K15013) and long-chain acyl-CoA synthetase (ACSL, K01897). At TP1, decreased abundance was observed in two ACSBG proteins in *A. hyacinthus* (FC = –0.60, *p*-value = 0.014; FC = –0.86, *p*-value = 0.043). At TP1, *P. lobata* also showed decreased abundance of ACSL (FC = –1.18, *p*-value = 0.019). At TP2, we observed a similar pattern in *A. hyacinthus*, with two ACSBG proteins showing decreased abundance (FC = –0.61, *p*-value = 0.033; FC = –1.02, *p*-value = 0.046), and ACSL also decreasing (FC = –0.93, *p*-value = 0.029). In contrast, at TP2, *P. lobata* showed increased abundance of ACSL (FC = 0.67, *p*-value = 0.000048).

#### Symbiosome maintenance proteins

We observed shifts in several key proteins potentially involved in symbiosomal maintenance. Two cathepsin proteins: CTSL (K01365) and CTSB (K01363), associated with the lysosome, were differentially abundant. At TP1, CTSB showed increased abundance in *A. hyacinthus* (FC = 0.79, *p*-value = 0.043) and *S. pistillata* (FC = 1.02, *p*-value = 0.015). CTSL also showed increased abundance in *A. hyacinthus* (FC = 1.59, *p*-value = 0.046) and *S. pistillata* (FC = 0.58, *p*-value = 0.025). No significant response was observed in *P. lobata* at TP1. At TP2, CTSB showed increased abundance in both *A. hyacinthus* (FC = 0.93, *p*-value = 0.028) and *P. lobata* (FC = 1.51, *p*-value = 0.028). CTSL showed a mixed response in *P. lobata*, with one copy increasing (FC = 1.50, *p*-value = 0.0015) and another decreasing (FC = –1.97, *p*-value = 0.022).

Niemann-Pick C2 (NPC2, K13443), a lysosome-associated protein, showed decreased abundance in *A. hyacinthus* at TP2 (FC = –2.04, *p*-value = 0.0069). Neutrophil cytosolic factor 2 (NCF2), involved in phagosomes and ROS production, showed decreased abundance at TP2 in both *A. hyacinthus* (FC = –0.88, *p*-value = 0.045) and *P. lobata* (FC = –1.63, *p*- value = 0.012). Carbonic anhydrase 2 (CA2, K18245) showed decreased abundance in *A. hyacinthus* at TP1 (FC = –2.26, *p*-value = 0.025) and TP2 (FC = –3.61, *p*-value = 0.0018).

Additionally, the transcription factor NF-κB (K02580) has been implicated to play a role in maintaining symbiosis (Mansfield et al., 2017), which showed increased abundance in *A. hyacinthus* at both TP1 (FC = 0.63, *p*-value = 0.025) and TP2 (FC = 1.43, *p*-value = 0.025). No significant shifts were observed in the other species [for additional signaling proteins see **Supplemental Text**]

### Tank effects in the endosymbiont proteome

After quality filtering of endosymbiont protein groups, we obtained 332, 275, and 184 proteins from the endosymbionts of *A. hyacinthus*, *P. lobata*, and *S. pistillata*, respectively. Protein abundances were subsequently normalized to account for changes in endosymbiont abundance across samples, thereby minimizing artifacts caused by bleaching-induced cell loss. PCA analysis, along with assessments of median protein intensities **[Supplemental File 2]**, revealed strong tank effects that led to proteomic shifts even among ambient samples, independent of thermal stress. These confounding effects limited our ability to interpret biological signal; thus, we did not pursue further analysis of the endosymbiont proteome.

### Differences in metabolomic biomarkers across species

We investigated metabolomic data to validate previous findings regarding dipeptides, amino acids, and antioxidants. The dipeptides KQ, RQ, RA and RV accumulate under thermal stress (Williams et al., 2021a). In *A. hyacinthus* at TP1, KQ (FC = 3.40, adj-*p*-value = 0.0049), RQ (FC = 2.30, adj-*p*-value = 0.026), and RV (FC = 1.86, adj-*p*-value = 0.0049) all significantly increased in abundance. At TP2, the same dipeptides remained elevated. In *S. pistillata* at TP1, only RV showed a significant increase (FC = 6.73, adj-*p*-value = 0.029), while the other dipeptides also had increased abundances, they were not statistically significant. In *P. lobata*, no dipeptides were significantly differentially abundant at either TP1 or TP2, though increased fold-changes were observed at TP2 **[Figure 4B, Table S6a]**.

**Figure 4.**
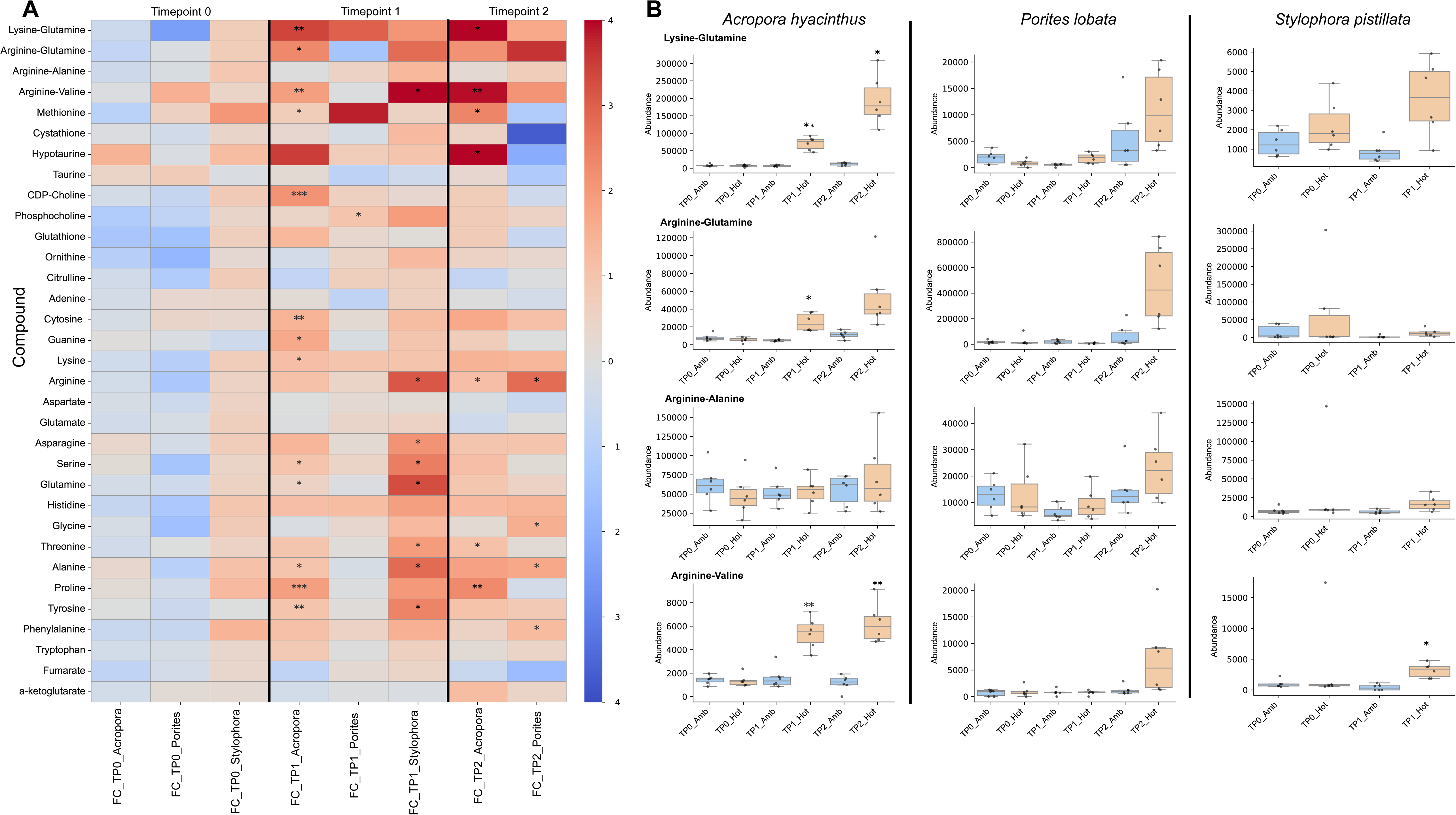
Differential abundance of polar metabolites. **(A)** Heatmap showing the various targeted metabolites in the study (y-axis) and the coral species and timepoints (x-axis). Red boxes indicate increased abundance, whereas blue boxes indicate decreased abundance. The asterisks denote statistical significance (* adj-*p*-value < 0.05; ** adj-*p*-value < 0.01; *** adj- *p*-value < 0.001). **(B)** Box plots for the four dipeptides assessed in the metabolomic dataset. Abundances (y-axis) are plotted for each dipeptide at each timepoint (x-axis). The thermally stressed samples are shown in orange, whereas ambient samples are shown in blue. Each column shows a different coral species and each row, a different dipeptide.

We assessed the abundance of well-characterized antioxidant metabolites (Williams et al., 2021b). Each coral holobiont showed a distinct profile of stress metabolite accumulation **[Table S6b]**. In *A. hyacinthus*, methionine had an increased abundance at both TP1 (FC = 0.74, adj-*p*-value = 0.046) and TP2 (FC = 2.34, adj-*p*-value = 0.018). Hypotaurine also had an increased abundance at TP2 in *A. hyacinthus* (FC = 11.23, adj-*p*-value = 0.035). CDP-choline also had an increased abundance in *A. hyacinthus* at TP1 (FC = 2.10, adj-*p*-value = 0.00030). However, these shifts were exclusive to *A. hyacinthus* only. Whereas phosphocholine was the only antioxidant metabolite that had an increased abundance at TP1 in *S. pistillata* (FC = 1.93, adj-*p*-value = 0.054) and a significant increase *P. lobata* (FC = 1.03, adj-*p*-value = 0.027) In addition, all three coral species showed increased abundance of amino acids **[Figure 4A, Table S6c]**. In *A. hyacinthus*, nine amino acids increased at TP1 (seven were statistically significant, adj-*p*-value < 0.05), and nine remained increased at TP2 (four were significant). In *S. pistillata*, 12 amino acids had an increased abundance (seven were significant). In *P. lobata*, seven amino acids had an increased abundance (four were significant). Of note only two amino acids (arginine and alanine) had an increased abundance in all three species, though this was temporally varied. Arginine had an increased abundance at TP1 in *S. pistillata* (FC = 3.01, adj-*p*-value = 0.035) and subsequently at TP2 in *A. hyacinthus* (FC = 1.12, adj-*p*-value = 0.036) and *P. lobata* (FC = 2.78, adj-*p*-value = 0.028). Alanine had an increased abundance at TP1 in *A. hyacinthus* (FC = 1.00, adj-*p*-value = 0.027) and *S. pistillata* (FC = 2.81, adj-*p*-value = 0.035), and at TP2 in *P. lobata* (FC = 1.69, adj-*p*-value = 0.017).

## Discussion

### Overall trends in coral physiology

The three sympatric coral species investigated in this study (*Acropora hyacinthus*, *Porites lobata*, and *Stylophora pistillata*) have well-documented differences in thermal stress response. *A. hyacinthus* is thermally sensitive, *P. lobata* is thermally resilient, and *S. pistillata* displays a low to moderate thermal stress tolerance (Burn et al., 2023; Fitt et al., 2009; Manalili et al., 2025; Meziere et al., 2022; Pratchett et al., 2020). In the experimental tanks, all three corals exhibited expected physiological responses to heat stress, including reduction in endosymbiont abundance (Glynn, 1991; Jokiel and Coles, 1977) and photosynthetic efficiency (*F_v_/F_m_*) (Warner et al., 1999, 1996). Consistent with the expected resilience levels, *A. hyacinthus* bleached first, wherein we observed a reduced endosymbiont abundance upon initial (TP1) stress and subsequently a more significant loss under prolonged (TP2) stress. Decline in photosynthetic efficiency was also observed. In contrast, *P. lobata* had only a slight decline (non-significant) in endosymbiont abundance, however, a significant decline in photosynthetic efficiency occurred under prolonged stress. Lastly, in *S. pistillata* colonies exhibited polyp bail-out, a rapid tissue loss response that is a severe stress reaction in this species (Chuang et al., 2021; Sammarco, 1982; Schweinsberg et al., 2021). Reduced photosynthetic efficiency was observed at initial stress. All three species hosted endosymbiotic algae from the genus *Cladocopium*, however the strains in each species were different. The three corals were collected from the same reef in the GBR, and likely had similar historical environmental stress profiles, helping to minimize variability due to differences in past stress exposure (Barshis et al., 2013; Kenkel and Matz, 2016; Morgan et al., 2017; Oliver and Palumbi, 2011; Risk and Edinger, 2011; Rogers et al., 2016; Safaie et al., 2018).

We assessed both the endosymbiont and host proteomes and observed tank effects in each. In the coral hosts, these effects were minimal, with only a few proteins showing differential abundance at TP0 (**Supplemental File 1**). In contrast, the endosymbiont proteome exhibited more pronounced variability. Endosymbiont protein abundances were normalized to account for bleaching-induced endosymbiont cell loss, however, this adjustment revealed exacerbated tank effects. Specifically, we observed a decline in protein abundance in ambient samples over time (**Supplemental File 2**), despite the absence of thermal stress. As a result, we excluded the endosymbiont proteome from further analysis and focused instead on host-specific responses to heat stress. Physiological indicators, including reductions in endosymbiont abundance and photosynthetic efficiency, supported the occurrence of bleaching. Together, these findings suggest a decline in photosynthate transfer from endosymbionts to the host under thermal stress, a response observed in previous coral bleaching studies (Davy et al., 2012; Grottoli et al., 2006).

### Pathway and protein level responses are consistent with diverged ecological outcomes to thermal stress

Corals depend on their algal symbionts to meet metabolic needs, thus, reduction in symbiont-derived photosynthate can severely impact holobiont fitness (Gordon and Leggat, 2010; Tremblay et al., 2012). Our data reveal that different coral species exhibit distinct proteomic responses to thermal stress and the associated decline in photosynthate transfer (**Figure 5**).

**Figure 5.**
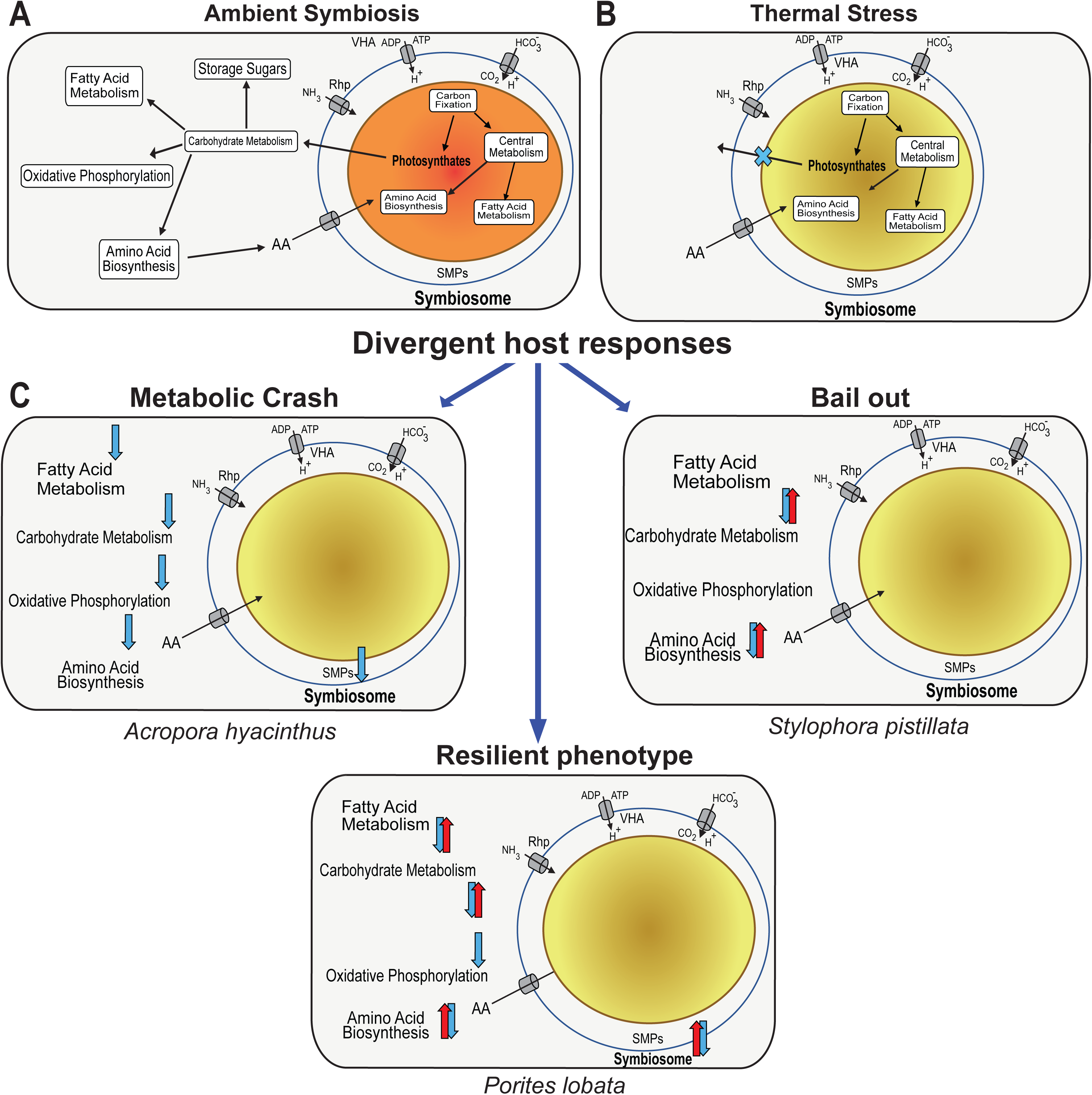
Divergent outcomes of thermal stress in three GBR coral species. (**A**) Diagram depicting a healthy symbiosis between the host and algal endosymbiont, involving transfer of nutrients from the host to the endosymbiont and subsequent transfer of photosynthates from the endosymbiont to the host. (**B**) A less efficient endosymbiont upon exposure to stress, limiting photosynthate transfer to host. The blue cross represents the lack of transfer of photosynthates from the endosymbiont to the host. (**C**) The divergent host metabolic responses of the three studied GBR coral species. In each panel, blue arrows represent pathways with proteins that decrease in abundance in response to thermal stress. The combination of a blue and red arrow represents a mixed protein response: i.e., a similar number of proteins in a pathway had increasing (red) and decreasing (blue) abundance.

However, the timing of these protein shifts was different between the species, likely due to differences in bleaching sensitivity, i.e., susceptible versus resilient. *A. hyacinthus* demonstrated early and extensive proteomic shifts, consistent with its high thermal sensitivity. At initial thermal stress, this species showed dramatic decreases in protein abundance across a substantial portion of its proteome (34.6% of proteins based on fold-change; **Supplemental Figure S3**). Upon prolonged thermal stress, there were signs of proteomic recovery. *P. lobata* displayed a more gradual and delayed response, wherein under initial stress a moderate response was observed. However, with prolonged stress, the response became more prominent, resembling the pattern seen in *A. hyacinthus*, but occurring later.

Notably, many proteins in *P. lobata* showed mixed responses across different copies, suggesting complex pathway regulation. *S. pistillata* exhibited a unique response pattern, showing both increases and decreases in protein abundances reflecting a mixed but active stress-response strategy, before ultimately undergoing polyp bail-out Consistent with their known ecological differences in thermal tolerance, we observed divergent responses at the pathway level among the three coral species. We evaluated the differential abundance of proteins across 25 KEGG pathways, 21 of which exhibited species-specific patterns. Among these, the “cell growth and death” pathway, which includes apoptosis and has been previously linked to coral thermal stress responses (Ainsworth et al., 2011), showed the most pronounced divergence and thus serves as a prime example [**Figure 3A**]. In *A. hyacinthus*, this pathway was downregulated during initial stress but upregulated under prolonged stress. In contrast, *P. lobata* exhibited a mixed initial response that shifted to downregulation under longer exposure to stress. *S. pistillata* also showed a mixed response at initial stress. These contrasting trajectories highlight the distinct stress-response strategies in these species and align with their known phenotypes. Such divergent responses underscore the challenges in identifying universal biomarkers of coral stress [**see below**].

Consistent with the known thermal sensitivity of species in the genus *Acropora* (Sakai et al., 2019; Singh et al., 2019), we observed that in *A. hyacinthus*, 16 pathways were downregulated under initial thermal stress, indicative of a broad metabolic collapse [**Figure 5**]. The host appeared to enter an energy-depleted state earlier than the other two coral species, putatively suggesting a higher reliance on endosymbiont derived photosynthates. We hypothesize that in response to reduced photosynthate transfer, the host shifts energy away from endosymbiont maintenance, reflected by decreased abundance of carbon supply proteins (e.g., CA2) (Bertucci et al., 2013; Zoccola et al., 2015) and increased abundance of proteins associated with symbiont expulsion (e.g., NF-κB) (Mansfield et al., 2017). These trends were reflected at the physiological level, wherein *A. hyacinthus* was the first of the three species to exhibit substantial endosymbiont loss. Under prolonged stress, there was an apparent recovery in the proteome, suggesting a compensatory shift towards mobilizing endogenous carbohydrate, lipid, and amino acid reserves to sustain metabolic function following bleaching (Rodrigues and Grottoli, 2007).

In *P. lobata*, a species known for its relative resilience to thermal stress (Huang et al., 2025; Rivera et al., 2022), few pathways were affected during initial stress. This finding was consistent with physiology, whereby no significant loss of endosymbiont cells was observed, and reduced photosynthetic efficiency was noted only under prolonged stress. Furthermore, under prolonged stress at an individual protein level, *P. lobata* had numerous instances of mixed responses (i.e., one copy of a protein had increased abundance and another showed a decrease). These mixed responses may indicate a compensatory mechanism in *P. lobata* which may be responsible for the higher thermal tolerance in this genus. Species in the genus *Porites* have larger nuclear genomes and subsequently encode more genes (30,000-60,000 genes) (Stephens et al., 2022), whereas other genera and species have smaller genomes and gene inventories, such as *Acropora* (∼20,000 genes) (Shinzato et al., 2021) and *Stylophora pistillata* (24,833 genes) (Voolstra et al., 2017). We speculate that the larger genomic repertoire in *Porites* species may enable a more complex and flexible stress response, potentially underpinning their thermal resilience phenotype.

Physiologically, *S. pistillata* exhibited a bail-out response under thermal stress (Schweinsberg et al., 2021). Unsurprisingly, at the molecular level we observed distinctive responses, compared to the other two species. We observed many pathways upregulated at initial stress (7 of the 25), which included transcription and translation, and were species-specific responses in *S. pistillata*. A study that performed salinity induced bail-out in the coral *Pocillopora acuta* found key enriched responses in signaling pathways (Chuang and Mitarai, 2020). Proteins in signaling pathways were relatively stable, however this may be due to the measurements being made prior to bail-out and not during active bail-out or may simply reflect the smaller number of proteins obtained from *S. pistillata* when compared to the other two coral species. Shared with *P. acuta*, we found an increased abundance in proteosome associated proteins that may be involved in proteolysis (PSMA3, PSMA4, and PSMD1) (Chuang and Mitarai, 2020) [**see Supplemental Text**].

### Diverged protein responses even in pathways with a conserved response

Only four of the 25 pathways exhibited a conserved response across all three coral species [**Figure 3A**]. Within these, two key trends emerged. First, although the direction of change was consistent across species (e.g., downregulation), the timing varied. For example, the energy metabolism pathway was downregulated in *A. hyacinthus* and *S. pistillata* during initial stress, but not until prolonged stress in *P. lobata* [**Figure 3A**]. These conserved, yet temporally staggered, responses align with known bleaching susceptibility, and subsequently endosymbiont photosynthate limitation. Declines in carbohydrate and energy metabolism have been widely documented, although the magnitude and timing differ across species and experimental conditions (Haydon et al., 2023; Petrou et al., 2021;). Lipid metabolism, a commonly used biomarker of coral bleaching, was also consistently downregulated, consistent with prior studies (Ermolenko and Sikorskaya, 2021; Grottoli et al., 2004; Imbs and Dembitsky, 2023). In contrast, protein folding, sorting, and degradation pathways were upregulated, which consist of heat-shock proteins, potentially reflecting increased demand for the turnover and clearance of misfolded or damaged proteins under stress (Petrou et al., 2021; Williams et al., 2021b).

The second trend we observed was that even within pathways exhibiting a conserved overall response, though temporally varied, the specific proteins driving that response differed across species. For instance, carbohydrate metabolism, which showed a consistent decrease across all three corals only 2 of the 39 differentially abundant proteins were shared between species. ALDH6A1 was differentially abundant in both *A. hyacinthus* and *P. lobata*, whereas ACO was affected in *S. pistillata* and *P. lobata*, but showed divergent expression patterns. The remaining 37 proteins were differentially abundant in only one species [**Figure 3B**].

### Shared protein and metabolite responses under thermal stress

#### Conserved protein responses

In this section, we examined the individual proteins that exhibited a conserved response in the three coral species. We first focused on canonical stress response proteins, commonly observed in corals and other metazoans. Heat shock proteins (HSPs) are well-established markers of thermal stress (Rosic et al., 2011; Sharp et al., 1997). These proteins have been widely used as biomarkers and play a central role in promoting survival under elevated temperatures in corals (Cleves et al., 2020; Louis et al., 2017). In our study, HSPA5 (also known as HSP70) and HSP90B showed conserved upregulation across all species. Notably, this response was temporally staggered wherein, *S. pistillata* exhibited increased abundance at initial stress, whereas *A. hyacinthus* and *P. lobata* showed increases later in the stress period. We also observed a conserved increase in calreticulin (CALR), a calcium-binding endoplasmic reticulum chaperone involved in protein folding and quality control (Fucikova et al., 2021). CALR has been proposed to act in concert with HSPs during thermal stress in corals (Ishibashi et al., 2023; Ruiz-Jones and Palumbi, 2017), supporting its role as a component of the conserved stress response machinery.

We also noted conserved proteins that are not directly involved in thermal stress responses but have been implicated in previous studies. One such protein is glutamine synthetase (GLNA), which is involved in ammonium assimilation. GLNA abundance decreased across all three corals; GLNA has been previously observed to have a decreased abundance at the transcriptomic level in *S. pistillata* from the Red Sea (Rädecker et al., 2021), suggesting a conserved response across species and geographic locations. Lastly, we observed a conserved response in two lysosomal cysteine proteases, cathepsin proteins CTSB and CTSL. Increased expression of CTSB under copper and thermal stress has been observed in corals and is suggested to play a role in the ROS response (Louis et al., 2017; Schwarz et al., 2013).

Increased CTSL expression has been previously associated with pathogenic infections and thermal stress in corals (Louis et al., 2017; Skutnik et al., 2020; Wright et al., 2015).

#### Conserved metabolomic responses

In this study, we employed polar metabolomics to identify and validate previously proposed metabolite biomarkers of thermal stress. Consistent with the physiological and proteomic responses, a notable temporal variation was observed in the metabolomic data. Within the coral holobiont, the host and endosymbiont share the same amino acid pool. Under thermal stress, amino acids serve two key functions: first, to maintain symbiosis, corals that transfer amino acids to their endosymbionts bleach more slowly than those that do not (Martinez et al., 2022b, 2022a, 2020); and second, to support host metabolism upon symbiont loss, as amino acid catabolism can compensate for reduced photosynthate supply (Rädecker et al., 2021). In this study, we observed an increase in free amino acid abundance in all three coral species, though the timing was species-specific. Only two amino acids, arginine and alanine, were consistently elevated across all species. Whereas the role of the increased amino acid pool remains unclear, species-specific physiological patterns suggest divergent strategies. In *A. hyacinthus*, increased amino acids coincided with early endosymbiont loss, suggesting they may supplement host metabolism under stress. Likewise, in *S. pistillata*, the increased amino acid pool, may be used to serve as a source of nutrition post bail-out. In contrast, *P. lobata* maintained stable endosymbiont abundances even under prolonged stress, implying that the elevated amino acid pool may be transferred to the symbiont to sustain the symbiosis.

Dipeptides are markers of stress in other model systems (Williams et al., 2021a) and show great promise in corals. Little is known however about the pathways involved in their synthesis and downstream functions. Stable isotope analysis demonstrates large differences in the turnover rates of dipeptides and free amino acids, suggesting that these have distinct biosynthetic mechanisms in corals (Chiles et al., 2022). We observed a conserved increase in dipeptide abundance across species, with all four dipeptides exhibited positive fold changes under thermal stress. Although only KQ, RQ, and RV were significantly increased in *A. hyacinthus*, and RV was significant in both *S. pistillata* and *P. lobata*. Like the patterns observed in the proteome, dipeptide responses were temporally staggered, reflecting species- specific dynamics.

### Study Limitations

Despite careful experimental design, tank effects were clearly observed in the coral and algal endosymbiont proteomes, which is a persistent issue in coral research (McLachlan et al., 2020; Molinari et al., 2025). After normalization of the endosymbiont proteome using changes in cell abundance, the magnitude of this effect became even more apparent (**Supplemental File 1**), which disallowed use of the algal data. Notably, colonies maintained under ambient, non-stress conditions still exhibited substantial shifts in endosymbiont proteomes across all three coral species, as shown by changes in median protein abundance. In contrast, tank effects on the coral host proteome were minimal. Whereas it is plausible that alterations in the endosymbiont proteome would influence the coral host, PCA and median protein abundance analyses suggest these downstream effects were not substantial (**Supplemental File 2**). In addition, we obtained different coverage levels in the proteome, wherein significantly fewer proteins were obtained in *S. pistillata* relative to the other species.

## Conclusion

In this study we subjected three sympatric coral species *A. hyacinthus* (sensitive), *P. lobata* (resilient), and *S. pistillata* (bail-out) with known ecological divergent responses to thermal stress. Consistent with the literature, we observed the expected physiological response for each species (i.e., reduced endosymbiont loss and decreased photosynthetic efficiency) in all three corals. However, we observe a clear diverged response in the proteome with only six pathways having a consistent response in all three corals. Even within the six pathways, we note that only a handful of proteins had a consistent response across species. We note that dipeptides consistently show an increased accumulation in all three species upon exposure to thermal stress, however, some species showed a stronger response than others, further supporting the species-specific nature of these responses. Overall, in *A. hyacinthus* we observe a sharp initial response to thermal stress followed by proteomic recovery, whereas in *P. lobata* we observe a mixed response only after prolonged stress, and lastly, in *S. pistillata* we observed an initial stress response which was followed by host bail-out.

## Methods

### Test species

A single large colony of *Acropora hyacinthus* (neat morphotype), *Stylophora pistillata*, and *Porites lobata* were sampled from Davies Reef (−18°49′30″S, 147°38′42″E) on the central GBR in February 2023 (Permit Number: G19:43148) (**Figure 1**). Only one genotype was used to avoid the possibility of stochastic inter-colony variation in the stress proteome and metabolome (Chille et al., 2024). These species were selected to represent different morphologies and heat stress sensitivities: *Acropora hyacinthus -* sensitive, tabulate, branching; *Stylophora pistillata –* low/moderate tolerance, branching; *Porites lobata -* tolerant, massive.

### Experimental design

Each colony was subdivided into 36 fragments (∼7cm^2^), mounted on experimental racks, and allowed to recover for 3 weeks prior to the start of the experiment at the National Sea Simulator (SeaSim) at the Australian Institute of Marine Science (AIMS) in outdoor flow through tanks replicating Davies Reef conditions (28.0°C, ca. ∼250 µmol photons m^-2^ s^-1^).

Prior to heat stress treatments, fragments were randomly assigned across six 50 L experimental flow through tanks (*n* = 18 per tank, 6 fragments per species), three for each of the two experimental treatments and acclimated for seven days before initiating heat stress. Fragments were exposed to two treatments: a temperature treatment of 32.4°C and a control maintained at 28.38°C (Maximum Monthly Mean at Davies Reef). The MMM is defined as the maximum sea surface temperature (SST) of the 12 monthly mean values for the years 1985-1990 and 1993 (Heron et al., 2014) temperature ramping increased from 28.38°C to 32.4°C, over 5 days (0.8°C/day) for treatment tanks. The heat stress experiment ran for a further 4 days concluding after 9 days. Heat exchangers and precision mixers were used to control tank water temperature within 0.1°C and programmed and monitored using the Supervisory Control and Data Acquisition system (SCADA) at SeaSim. Experimental tanks were immersed within temperature controlled exterior insulating jackets of seawater kept at treatment temperatures (one jacket per treatment) and each tank fitted with a TC Direct PT- 100 temperature probe to maintain ramp rate and programmed temperatures. Filtered seawater was continuously supplied to the experimental system at an inflow rate of 0.8L/minute and each tank was fitted with a circulator (Turbelle nanostream 6015 3.5 W, Tunze Aquarientechnik Gmbh, Germany) for water movement. Light was on a 24 hour 12:12 light/dark cycle with a 2-hour sunrise/sunset ramp time (half-sine ramp) and maximum photosynthetically active radiation (PAR) at 280 µmol/m^2^/s supplied from SS Gen 1 Customised multichip LEDs (325 W, The National Sea Simulator, Townsville, Australia) and measured continuously in each tank using a Li-192 sensor (UWQ8687, Li-COR, USA) and Li-250Am meter (Li-COR, USA). Water temperature and PAR was stable at target levels throughout the experiment.

### Sampling plan

Corals were sampled at three time points **(Figure 1**): TP0, day 0, at the start of the experiment (end of coral acclimation described above) but prior to temperature ramp-up; TP1, day 5, at the end of temperature ramp-up; and TP2, day 9, four days after the end of temperature ramp-up. At each timepoint, sampling included photographs of each fragment [**Supplemental Figure 1**] with one fragment per species from each tank (n = 3/treatment/species) immediately snap frozen and kept at -80°C until processing for metabolomics, proteomics, and algal endosymbiont cell counts and typing (see below). At each timepoint, and additionally, on days 7 and 8, the maximal quantum yield (*F_v_/F_m_*) of chlorophyll a fluorescence of all fragments was measured using a Diving-PAM (Waltz, Germany; Fiberoptics outer diameter 8 mm; settings: measuring light intensity = 6, saturation pulse intensity = 10, saturating width = 0.8s, damping = 2, gain = 4). Measurements were made before dawn therefore, colonies were low light acclimated (Suggett et al., 2022). Each fragment was measured three times, at separate parts of the fragment, and the average values were recorded. Analysis of variance (ANOVA) was utilized to assess the difference in *F_v_/F_m_*for each coral species. Additionally, Tukey-HSD test was performed to do a pairwise comparison between ambient and thermal stress at each timepoint.

### Endosymbiont counts

A small section of tissue (∼1 cm^2^) from each frozen fragment collected at each timepoint (TP0, TP1, and TP2), was air-brushed in PBS buffer, separated (as above), and washed three times in filtered seawater, these cells were used to calculate endosymbiont abundance. Cell quantification was determined visually *via* a haemocytometer (Neubauer Haemocytometer, Fisher Scientific, Loughborough, UK) with eight replicate quadrats counted for each sample. Surface area measurements of each air-brushed coral skeleton were conducted using the single wax dipping method (Veal et al., 2010). Eight cylindrical calibration standards were measured using digital callipers with a precision of 0.001 mm to determine their geometric surface areas. These were used to generate a standard curve relating wax mass to surface area. Coral skeletons were weighed and then skeletons and calibration standards were immersed in paraffin wax at 65L°C for 2 seconds, then removed and gently rotated to promote uniform coating and initial air drying. Samples were subsequently left to dry for an additional 15 minutes at ambient temperature before being reweighed. The surface area of each skeleton was then calculated based on the difference in mass before and after wax coating, using the calibration curve.

Endosymbiont cell densities were subsequently normalized using surface area, i.e., cell density / surface area (cells/ml/cm^2^), hereinafter cell abundance. Normalized cell abundance were log_2_ transformed for visualization. Analysis of variance (ANOVA) was utilized to assess the difference in normalized cell density for each coral species. Additionally, Tukey-HSD test was performed to do a pairwise comparison between ambient and thermal stress at each timepoint.

### Endosymbiont typing

A small section of tissue (∼2 cm^2^) from each frozen fragment collected at TP0 was air- brushed in PBS buffer for endosymbiont typing using amplicon sequencing of the ribosomal Internal Transcribed Spacer 2 (ITS-2) region. The tissue slurry was centrifuged at 3000g for 10 minutes to separate the coral host and algal symbiont fractions. DNA was extracted from the algal pellet using the Qiagen DNeasy Blood and Tissue kit (Qiagen, Germany) following the “protocol: purification of total DNA from animal tissues” including a RNase A-based RNA digestion procedure. Genomic DNA was sent to the Ramaciotti Centre for Genomics (University of New South Wales, Sydney, Australia) and ITS-2 amplicons sequenced using the sym_var_5.8s2/sym_var_rev primers on the NextSeq 1000 P1 platform with 2x300bp paired-end sequencing (including PhiX spike-in) (Hume et al., 2018). ITS2 sequences were submitted to the SymPortal analytical framework for quality control and ITS2 type profile analysis (https://symportal.org) as described by Hume and coauthors (Hume et al., 2019).

### Protein extraction

Proteins were extracted using a RIPA lysis buffer, composed of Tris-HCl (50mM), NaCl (150mM), SDS (1%), and DI water. The buffer was chilled on ice until ready for use and one cOmplete mini EDTA-free tablet was dissolved in 10 ml of lysis buffer immediately before extraction. A total of 500μl of ice-cold RIPA lysis buffer was added to a tube filled with 0.5mm silica beads. Approximately 0.2-0.4 g of coral sample was clipped into each bead tube. The samples were vortexed for at least five minutes, or until the skeleton was stripped from the tissue. The samples were then incubated on ice for 30 minutes, before centrifuging at 10,000 rcf for 10 minutes and extracting the supernatant. Protein quality was assessed using a Qubit protein assay. Samples were stored in a freezer at -80°C.

### Proteomics analysis

Proteomic analysis was performed at the University of South Florida Proteomics Core. The proteins were normalized before digestion using 25μg for each sample. After digestion, an equal amount of sample (1000ng) was injected for each run. LC-MS analysis peptides were characterized using a Thermo Q-exactive-HF-X mass spectrometer coupled to a Thermo Easy nLC 1200. Samples separated at 300nl/min on an Acclaim PEPMAP 100 trap (75uM, 2CM, c18 3um, 100A) and an Thermo easy spray column (75um, 25cm, c18, 100A) using a 180- minute gradient with an initial starting condition of 2% B buffer (0.1% formic acid in 90% Acetonitrile) and 98% A buffer (0.1% formic acid in water). Buffer B was increased to 28% over 140 minutes, then up to 40% in an additional 10 minutes. High B (90%) was run for 15 minutes afterwards. The mass spectrometer was outfitted with a Thermo nanospray easy source with the following parameters: Spray voltage: 2.00V, Capillary temperature: 300dC, Funnel RF level=40. Parameters for data acquisition were as follows: for MS data the resolution was 60,000 with an AGC target of 3e^6^ and a max IT time of 50 ms, the range was set to 400-1600 m/z. MS/MS data was acquired with a resolution of 15,000, an AGC of 1e^4^, max IT of 50 ms, and the top 30 peaks were picked with an isolation window of 1.6m/z with a dynamic execution of 25s.

For each of the three species, a comprehensive metaproteomic database was assembled based on the ITS2 results (**Figure 2C**) to aid protein identification. A proteomic database for *Porites lobata* was assembled from: (1) predicted genes from the *P. lobata* transcriptome assembly (v1) (Bhattacharya et al., 2016); (2) predicted genes from the *P. lobata* (KU572435) mitochondrial genome assembly (Tisthammer et al., 2016); (3) predicted genes from the *Cladocopium goreaui* nuclear genome assembly (Chen et al., 2022), (4) predicted genes from the *Breviolum minutum* mitochondrial genome assembly (Shoguchi et al., 2015); and (5) predicted genes from the *Cladocopium* C3 plastid genome assembly (Barbrook et al., 2014). A proteomic database for *Acropora hyacinthus* was assembled from: (1) predicted genes from the *A. hyacinthus* nuclear genome assembly (v1) (Singh et al., 2019); (2) predicted genes from the *A. hyacinthus* (OP311657) mitochondrial genome assembly (Wang et al., 2022); (3) predicted genes from the *Cladocopium goreaui* nuclear genome assembly; (4) predicted genes from the *Breviolum minutum* mitochondrial genome assembly; and (5) predicted genes from the *Cladocopium* C3 plastid genome assembly. Finally, a proteomic database for *Stylophora pistillata* was assembled from: (1) predicted genes from the *S. pistillata* nuclear genome assembly (v1) (Voolstra et al., 2017); (2) predicted genes from the *S. pistillata* (EU400214) mitochondrial genome assembly; (3) predicted genes from the *Cladocopium goreaui* nuclear genome assembly; (4) predicted genes from the *Breviolum minutum* mitochondrial genome assembly; and (5) predicted genes from the *Cladocopium* C3 plastid genome assembly.

### Protein clean-up and normalization

Raw proteomic data was generated using Max quant 2.0.3.1 (Cox and Mann, 2008; Tyanova et al., 2016). Proteomic data cleaning was conducted in RStudio v2024.04.2+764 using R v4.4.1 and the protti R package v0.9.0 (Quast et al., 2022). This package was used to clean, normalize, and generate pre- and post-cleaning reports for each of the three host species.

Assessments of peak intensity distribution, data completeness, coefficient of variation across sites, and principal component analysis (PCA) were performed before and after cleaning. To create a high-quality protein group library, LC-MS results were filtered to exclude symbiont proteins, proteins with greater than one unique peptide and razor peptide for each protein group (Unique peptides > 1 & Razor + unique peptides > 1) and a protein group Q-value less than 0.01. Proteins belonging to a set of potential contaminants provided by the LC-MS facility were also removed. The data matrix was then filtered to exclude observations with intensity values below 5, which likely represent false assignments. Protein intensities were log2-transformed using the dplyr *mutate* and base R *log2* functions. Outlier samples were identified by PCA and data completeness using the protti *qc_pca* and *qc_data_completeness* functions. Based on these analyses, two samples with low completeness and that were PCA outliers in their respective datasets were removed, Acropora_638 and Stylophora_2312.

Protein groups missing from more than 80% of samples were also removed. This threshold allowed proteins present in at least 80% of samples from a species to remain in the dataset, therefore it had to have been missing in >3 samples to be removed. For the remaining high-confidence observations, missing values were imputed using the *missForest* function from the missForest package with default parameters (maxiter = 10, ntree = 100) and a set seed of 124. Finally, log2-transformed peak intensities were median-normalized per treatment_timepoint group using the protti *normalise* function, resulting in the final cleaned protein abundance matrix.

### Downstream proteomic analysis

GhostKOALA (v2.0) (Kanehisa et al., 2016) was used to annotate proteins with KEGG Orthology (KO) numbers. We subsequently calculated the log_2_ Fold-Change (hereinafter, FC) for each protein between the ambient and high temperature samples for TP0, TP1, and TP2 using Python (v.3.9.12) (additional packages pandas v1.4.2 and numpy v1.26.4), and using SciPy (v1.12.0) to assess statistical significance with a student t-test (*n* = 3). A protein was considered to have an increased abundance if it had a FC > 0.5 and *p*-value < 0.05, likewise, a protein was considered to have a decreased abundance if it had a FC < -0.5 and *p*-value < 0.05. However, upon correcting for multiple testing (Benjamini Hochberg corrections), no adjusted *p* – value was < 0.05, potentially due to the high variation in the proteomic data. We therefore proceed with only *p-*value based analyses. We also performed a pathway level analysis, using the “B-level Description” annotation generated from the KEGG database. A pathway was upregulated if 5% of the total proteins in a pathway had an increased abundance and was downregulated if 5% of the total proteins had a decreased abundance. A pathway was considered to have a mixed response if both upregulation and downregulation was observed.

### Metabolomic analysis

In addition to physiological and proteomic analyses, we generated untargeted polar metabolomics data to assess metabolite abundance (notably dipeptides) previously linked to thermal stress responses in corals (Williams et al., 2021a). Metabolites were extracted from preserved coral tissue using a 40:40:20 methanol:acetonitrile:water buffer with 0.1 M formic acid, followed by homogenization, centrifugation, acid neutralization with ammonium bicarbonate, and transfer to autosampler vials for LC-MS analysis. LC-MS analysis was performed at the Metabolomics Core Facility at the Cancer Institute of New Jersey. Prior to sample runs, LC-MS performance was verified using commercial and in-house standards to ensure mass accuracy and signal consistency. Samples were randomized to minimize batch effects, and method blanks were included to assess background signals. Metabolite separation was performed using HILIC on a Vanquish UHPLC (Thermo Fisher Scientific, Waltham, MA) and the subsequent full scan mass spectrometry was performed on the Thermo Q Exactive PLUS mass spectrometer operating in both positive and negative modes. Data were processed using MAVEN (Melamud et al., 2010) with identification based on accurate mass and retention time matching to an in-house library, and low-abundance metabolites were filtered from downstream analyses. For an in-depth description of the extraction protocols and mass-spectrometry settings **see supplemental text.**

We subsequently calculated the log_2_FC for each of the metabolites across the species, comparing the ambient and thermally stressed treatments at each timepoint. We also assessed the statistical significance for each of the metabolites using a Welch t-test (*n* = 6) with multiple testing correction using the Benjamini-Hochberg procedure, using SciPy (v1.12.0).

## Acknowledgments

We acknowledge the Bindal and Wulgurukaba Peoples as the Traditional Custodians of the land and sea Countries where the Great Barrier Reef part of this research took place. We wish to acknowledge Elders past, present, and emerging, and their continuing spiritual connection to sea Country. Thank you to the staff of the AIMS vessel and the National Sea Simulator for field and laboratory support. We also thank Elizabeth Wood at the University of South Florida, Proteomics Core for support with proteomic data collection and analysis. We thank Eric Chiles and Dr. Xiaoyang Su at the Rutgers Metabolomic Core. Finally, we wish to thank Eric Hochberg from the Bermuda Institute of Ocean Sciences for his contributions to the design of the study and assistance with collection of the samples.

## Author contributions

Conceptualization: SN, TGS, EEC, SG, LKB, DB

Methodology: SN, TGS, EEC, SG, LKB, DB

Investigation: SN, TGS, EEC, SG, LKB, DB

Formal Analysis: SN, TGS, EEC

Supervision: LKB, DB

Writing (original draft): SN, TGS

Writing (review & editing): SN, TGS, EEC, SKG, LB, DB

## Data and materials availability

All data are available in the main text or the supplementary materials. The GBR coral mass spectrometry proteomics data have been deposited to the ProteomeXchange Consortium *via* the PRIDE partner repository with the dataset identifier PXD052478 (https://www.ebi.ac.uk/pride/). The metabolomic mass spectrometry data are available from the MassIVE database with the dataset identifier MSV000094832 (https://massive.ucsd.edu/). The ITS2 sequence read files are available from the Sequencing Read Archive (SRA) with the dataset identifier PRJNA1119670 (https://www.ncbi.nlm.nih.gov/sra/).

## Supplemental Information

Table S1a. Endosymbiont cell density and photosynthetic efficiency for samples from all three GBR coral species.

Table S1b. SymPortal ITS2 Symbiodiniaceae communities within Acropora hyacinthus, Porites lobata and Stylophora pistillata from the Great Barrier Reef. ITS2 sequence data and major ITS2- type profiles.

Table S2. Proteomic data for the Acropora hyacinthus host across timepoints, with differential accumulation statistics and KEGG annotations shown for each protein.

Table S3. Proteomic data for the Porites lobata host across timepoints, with differential accumulation statistics and KEGG annotations shown for each protein.

Table S4. Proteomic data for the Stylophora pistillata host across timepoints, with differential accumulation statistics and KEGG annotations shown for each protein.

Table S5. Summary table of all GBR coral host proteins detected for each species across the stress experiment for specific pathways. Each protein is counted only once here for reference. Multiple copies of a given protein are included in these counts.

Table S6a. Dipeptide metabolite abundance variation between control and treatment conditions at each time point from the GBR corals.

Table S6b. Annotated antioxidant metabolite abundance variation between control and treatment conditions at each time point from the GBR corals.

Table S6c. Amino acid metabolite abundance variation between control and treatment conditions at each time point from the GBR corals.

Supplemental File 1: Endosymbiont proteome distribution after cell density normalization and principal component analysis. Box and whisker plots of log2 protein intensities across treatments and timepoints for A. hyacinthus (A), P. lobata (C), and S. pistillata (E) symbiont proteins. Protein intensities were group-median normalized by treatment_timepoint for each species independently to examine protein-wide up- or down- regulation in response to treatment. PCA plots of the first two principal components based on group-normalized log2 protein intensities for A. hyacinthus (B), P. lobata (D), and S. pistillata (F). Legends describing the colors and shapes are shown to the right of each figure.

Supplemental File 2: Host protein intensity distribution and principal component analysis. Box and whisker plots of log2 protein intensities across treatments and timepoints for A. hyacinthus (A), P. lobata (C), and S. pistillata (E) host proteins. Protein intensities were group-median normalized by treatment_timepoint for each species independently to examine protein-wide up- or down- regulation in response to treatment. PCA plots of the first two principal components based on group-normalized log2 protein intensities for A. hyacinthus (B), P. lobata (D), and S. pistillata (F). Legends describing the colors and shapes are shown to the right of each figure.

Supplemental Figure 1: Coral fragment images over the course of the experiment. This figure shows coral fragments from all three species at various timepoints during the experiment. Timepoints and treatment conditions are indicated on the left, and fragment IDs are shown below each image. In Stylophora pistillata, distinct paling of the host colonies is visible as early as TP1 under thermal stress. Similarly, Acropora hyacinthus displays clear signs of bleaching by TP2 based on visual assessment. In Porites lobata, mild paling is also evident by TP2.

Supplemental Figure 2: Host total proteome distribution. Volcano plot of the proteins identified in the host proteome data from the three sympatric species. The plots are arranged horizontally by species and vertically by time point. Data is not available for S. pistillata TP2 due to the bail out phenotype, and as such, the respective position are filled with “NA”. The dots in each plot represent individual proteins, with the blue dots representing proteins with a statistically significant decreased abundance (FC < -0.5, p-value < 0.05) and red dots representing proteins with a statistically significant increased abundance (FC > 0.5, p-value < 0.05) at the given time point. A legend describing the colors is shown at the bottom of the figure.

Supplemental Figure 3: Fold-change based pathway level assessment. (A) Stacked mirror plot of differentially abundant proteins at TP1 in each of the three colonies. “B-Description” (KEGG classification) is presented on the x-axis and the number of proteins that are differentially abundant on the y-axis. Note only pathways with more than 5 proteins differentially abundant (across all species) have been included in the plot. The different species have been shown in different colors. Bars extending up are the number of proteins that have an increased abundance and down show decreased abundance. (B) Similar to (A) but shows differentially abundant proteins at TP2. Data is only available for A. hyacinthus and P. lobata.

