## Supplemental Text for "Metaproteome analysis of short-term thermal stress in three sympatric coral species reveals divergent host responses"

**Methods**

**Metabolomic workflow**

In addition to the physiological measurements and proteomic work we also generated untargeted polar metabolomics data from the three corals used in this study. Our goal was to assess the abundance of metabolites (most notably, dipeptides), whose increasing abundance has previously been shown to be associated with the thermal stress response in *M. capitata* and other Hawaiian coral species (Williams et al., 2021). Metabolites were extracted using a protocol optimized for water-soluble polar metabolite analysis on LC-MS. The extraction buffer was a solution of 40:40:20 (methanol:acetonitrile:water) (v/v/v) + 0.1 M formic acid. The extraction buffer was stored at −20°C before usage. Immediately preceding the metabolite extraction, 1 ml of extraction buffer was added to the 2ml tube with glass beads. Pieces (~0.2-0.4 g) of the -80°C preserved nubbins were then clipped, weighed, and added. The coral samples were then homogenized. The rest of the homogenate was then transferred to a 1.5-ml Eppendorf tube, vortexed for 10 s, and then centrifuged for 10 min at 14,000*g* at 4°C. After centrifugation, there was a pellet at the bottom of the tube. A final 500 μl aliquot of the homogenate was then pipetted to a second clean Eppendorf tube, to which 44 μl of 15% NH_4_HCO_3_ was added to neutralize the acid in the buffer. This was the final extract and was transferred to autosampler vials to be loaded to instrument.

Prior to running the samples, the LC-MS system was evaluated for performance readiness by running a commercially available standard mixture and an in-house standard mixture to assess the mass accuracy, signal intensities and the retention time consistency. All known metabolites in the mixture were detected to within 5ppm mass accuracy. Method blank samples matching the composition of the extraction solvent were used in every sample batch to assess background signals and ensure there was no carryover from one run to the next. In addition, the sample queue was randomized with respect to sample treatment to eliminate the potential for batch effects.

The HILIC separation was performed on a Vanquish Horizon UHPLC system (Thermo Fisher Scientific, Waltham, MA) with XBridge BEH Amide column (150 mm × 2.1 mm, 2.5 μm particle size, Waters, Milford, MA) using a gradient of solvent A (95%:5% H 2 O:acetonitrile with 20 mM acetic acid, 40 mM ammonium hydroxide, pH 9.4), and solvent B (20%:80% H 2 O:acetonitrile with 20 mM acetic acid, 40 mM ammonium hydroxide, pH 9.4). The gradient was 0 min, 100% B; 3 min, 100% B; 3.2 min, 90% B; 6.2 min, 90% B; 6.5 min, 80% B; 10.5 min, 80% B; 10.7 min, 70% B; 13.5 min, 70% B; 13.7 min, 45% B; 16 min, 45% B; 16.5 min, 100% B and 22 min, 100% B (Su et al., 2020). The flow rate was 300 μl/min. Injection volume was 5 μL and column temperature was 25 °C. The autosampler temperature was set to 4 °C and the injection volume was 5µL. The full scan mass spectrometry analysis was performed on a Thermo Q Exactive PLUS with a HESI source which was set to a spray voltage of -2.7kV under negative mode and 3.5kV under positive mode. The sheath, auxiliary, and sweep gas flow rates of 40, 10, and 2 (arbitrary unit) respectively. The capillary temperature was set to 300 °C and aux gas heater was 360 °C. The S-lens RF level was 45. The m/z scan range was set to 72 to 1000 m/z under both positive and negative ionization mode. The AGC target was set to 3e6 and the maximum IT was 200ms. The resolution was set to 70,000.

The full scan data were processed with a targeted data pipeline using MAVEN software package (Melamud et al., 2010). Compound identification was assessed using accurate mass and retention time match to the metabolite standards from the in-house library. Metabolites in the targeted dataset were filtered for low abundance: i.e., if more than 3 replicates had no detected intensity, they were excluded from downstream analysis.

**Results**

*Signal Transduction*

We also assessed MAPK2 (K04368), part of the MAPK signaling pathway, which showed decreased abundance in A. hyacinthus (FC = –0.95, p-value = 0.016) and P. lobata (FC = –0.72, p-value = 0.0085) at TP1 only. A suite of significant shifts in ATP-based transporters was also observed. In *A. hyacinthus* at TP1, we observed a decreased abundance of ATP1A (FC = -0.62, *p*-value = 0.014) and two copies of ATP2B (FC = -0.55, *p*-value = 0.011and FC = -1.09, *p*-value = 0.047). In *P. lobata* at TP1, we observe that ATP2B (FC = -0.59, *p*-value = 0.00072) and ATP2A (FC = -0.53, *p*-value = 0.033) have decreased abundance. In *S. pistillata* at TP1, we observed two copies of ATP1A, both with decreased abundance (FC = -0.54, *p*-value = 0.020 and FC = -0.81, *p*-value = 0.019). Another ATP transporter, ATP2A had an increased abundance (FC = 0.70, *p*-value = 0.0099). Furthermore, at TP2 we observe no significant shifts in these proteins in *A. hyacinthus*, and a mixed response was observed in *P. lobata*, i.e., ATP2B (FC = -0.79, *p*-value = 0.0076) and in ATP2A (FC = 0.67, *p*-value = 0.019).

*Protein folding, sorting, and degradation*

Proteasome associated proteins, including PSMA3 (K02727), PSMA4 (K02728), PSMA7 (K02731), PSMB1 (K02732), PSMD1 (K03032), PSMD7 (K03038), PSMD8 (K03031), have diverged responses across the three coral species. At *S. pistillata* at TP1, PSMA3 (FC = 0.50, *p*-value = 0.038), PSMA4 (FC = 0.55, *p*-value = 0.047), and PSMD1 (FC = 0.55, *p*-value = 0.024) all had an increased abundance. Whereas, in *P. lobata* at TP1, a decreased abundance in the following proteins was observed, PSMA7 (FC = -0.50, *p*-value = 0.044), PSMD7 (FC = -1.16, *p*-value = 0.043), and PSMD8 (FC = -1.16, *p*-value = 0.020). In *A. hyacinthus* at TP1*,* a decreased abundance is observed only in the PSMB1 protein (FC = -0.96, *p*-value = 0.030). Furthermore, in *P. lobata* at TP2, a similar trend was observed in the proteins PSMD2 (FC = -0.71, *p*-value = 0.031), PSMD3 (FC = -2.07, *p*-value = 0.047), and PSMD7 (FC = -0.76, *p*-value = 0.046). Lastly, in *A. hyacinthus* at TP2, a decreased abundance was also observed in PSMA6 (FC = -2.15, *p*-value = 0.042).

***Enzymatic ROS response***
We observed six proteins associated with ROS detoxification: SOD1 (K04565), SOD2 (K04564), catalase (CAT, K03781), glutathione peroxidase (GPX, K00432), glutathione reductase (GSR, K00383), and glutathione synthetase (GSS, K21456). Only GSS showed a significant decrease at TP1 in P. lobata (FC = –0.56, p-value = 0.022). Aside from this, overall levels of these enzymes remained stable throughout the experiment. In A. hyacinthus at TP1, SOD1 showed decreased abundance (FC = –0.51, p-value = 0.43), returning to ambient levels at TP2. One GPX copy decreased at both TP1 (FC = –0.90, p-value = 0.25) and TP2 (FC = –0.82, p-value = 0.12). One GSR protein decreased at TP1 (FC = –0.53, p-value = 0.14), again returning to baseline at TP2. In P. lobata, shifts were observed only at TP2: one CAT protein increased (FC = 1.43, p-value = 0.11), while one GPX protein decreased (FC = –0.67, p-value = 0.13). No significant changes were observed in S. pistillata.
